# Structure of *Geobacillus stearothermophilus* Cas9: insights into the catalytic process and thermostability of CRISPR-Cas9

**DOI:** 10.1101/2024.06.05.595678

**Authors:** Panpan Shen, Lilan Zhang, Beibei Liu, Xian Li, Jian Min, Jian-Wen Huang, Chun-Chi Chen, Rey-Ting Guo

**Affiliations:** State Key Laboratory of Biocatalysis and Enzyme Engineering, Hubei Hongshan Laboratory, Hubei Collaborative Innovation Center for Green Transformation of Bio-Resources, Hubei Key Laboratory of Industrial Biotechnology, School of Life Sciences, Hubei University, Wuhan 430062, PR China; Department of Immunology and Pathogen Biology, School of Basic Medical Sciences, Hangzhou Normal University, Hangzhou 311121, China

**Keywords:** CRISPR-Cas9, cryo-electron microscopy, *Geobacillus stearothermophilus*, thermostability, catalytic mechanism, L1-crevice

## Abstract

CRISPR-Cas9 has been developed as a powerful gene editing tool, but the mechanism governing the intricate catalytic process remains incompletely resolved. Here, the cryo-electron microscopy structures of thermostable Cas9 from *Geobacillus stearothermophilus* (GeoCas9) in complex with sgRNA and target DNA are reported. The structure of GeoCas9 in complex with sgRNA reveals a slit termed L1-crevice comprising HNH, RuvC, and L1 helix as a transient storage site of 5’ spacer of sgRNA. When 5’ spacer is extracted to pair with the target DNA, L1-crevice collapses to trigger the subsequent HNH domain translocation. In addition, structural and biochemical analyses suggest that the resilience of GeoCas9 at elevated temperature is related to the unique PI domain conformation. These results advance our understanding into the catalytic process of Cas9 and unveil the molecular mechanism that accounts for the superior thermal profile of GeoCas9.

## Introduction

Clustered regularly interspaced short palindromic repeats (CRISPR) is an RNA-guided adaptive immunity in bacteria and archaea that is exploited to defend against invading plasmids and phages^1,2^. In conjugation with CRISPR-associated (Cas) proteins, this machinery can recognize and cleave nucleic acids that are complementary to the CRISPR sequence. Therefore, the CRISPR-Cas systems have been developed as promising gene editing tools that hold great potentials in various biotechnological areas including diagnostic and therapeutic applications^3,4^.

The most common and best-studied CRISPR-Cas system is CRISPR-Cas9, which carries a guide RNA to cleave the target double stranded DNA (dsDNA) that contains a complementary sequence and Protospacer Adjacent Motif (PAM). A functional CRISPR-Cas9 system comprises a single effector multi-modular Cas9 protein that contains HNH (His-Asn-His) endonuclease domain and RuvC-like endonuclease domain to cleave target-(TS-DNA) and non-target strand DNA (NTS-DNA), respectively^5^. The second component is CRISPR transcribed RNA (crRNA) that comprises a short segment known as spacer (17-24 nt in length) on the 5’-end that is complementary to TS-DNA and a repeat sequence on the 3’-end. The third part is trans-activating crRNA (tracrRNA) that base pairs with the repeat sequence of crRNA and plays a role in recruiting Cas9^6^. These two RNAs are fused to yield a single guide RNA (sgRNA) to simplify the application of CRISPR-Cas9^7^. By designing the sequence of the spacer, the CRISPR-Cas9 system can be programmed to target any given DNA sequence of interest^8^.

The catalytic process of CRISPR-Cas9 has been illustrated via structural, biochemical and computational approaches^9,10^. This process begins from the loading of sgRNA to the apo-form Cas9 to afford a binary ribonucleoprotein complex. The target dsDNA that carries the spacer-complement protospacer and cognate PAM binds to the PAM-recognition site of Cas9 and is unwound to TS- and NTS-DNA. The NTS-DNA is cleaved by the RuvC domain. The TS-DNA pairs with the spacer region of the sgRNA to form a RNA:DNA duplex. Then the HNH domain along with the L1 helix on its N-terminus conduct dramatic conformation change to approach the protospacer site on the RNA:DNA duplex to cleave the TS-DNA^8^. The domain transition has been well-illustrated by a number of structures of Cas9 in complex with sgRNA or target DNA^11–19^. Compared with the wealth information of the substrate DNA-bound state, there are only four Cas9 complex structures containing sgRNA from *Streptococcus pyogenes*, *Neisseria meningitides*, *Hemophilia parainfluenza* and *Staphylococcus aureus*^11,12,20,21^, with the latter two containing specific inhibitors, which might limit the comprehensive understanding of the catalytic process of Cas9.

CRISPR-Cas9 systems can be divided into subtype A, B and C depending on the inclusion of alternative associated Cas proteins and the degree of homology between Cas9 proteins^22,23^. The well-studied SpyCas9 from *S. pyogenes* belongs to type A^24^, and very few studies about type B have been reported that only accounts for around 3% Cas9^22^. Type C system is the simplest and accounts for around half of reported Cas9 including Nme1Cas9 from *N. meningitidis*^12^, CjeCas9 from *Campylobacter jejuni*^13^, CdiCas9 from *Corynebacterium diphtheriae*^17^, AnaCas9 from *Actinomyces naeslundii*^25^, AceCas9 from *Acidothermus cellulolyticus*^16^, HpaCas9 from *H. parainfluenzae*^11^ and GeoCas9 from *Geobacillus stearothermophilus*^26^. Notably, many of them are from extremophiles that habitat at elevated temperature or acidic/alkaline pH, which are considered attractive gene editing tools owing to their stability under extreme conditions^23,27,28^.

GeoCas9 that exhibits the optimal activity at 50-60 □ and can operate at 75 □ is among the most thermophilic Cas9 reported so far^26^. The outstanding thermostability renders GeoCas9 an attractive gene editing tool at elevated temperature as well as in thermophilic organisms. A recent study has reported the crystal structure of HNH domain of GeoCas9, which suggests a residue that correlates with the domain stability but shows minimal effect to the behavior of the full-length Cas9^29,30^. It is suspected that the overall architecture of the ribonucleoprotein complex should be required to investigate the mechanism underlying the unique thermal profile of GeoCas9. In this study, we solved GeoCas9 structure in complex with sgRNA with or without target dsDNA by using single particle cryo-electron microscopy (cryo-EM). These results not only increase our understanding of type C Cas9, but reveal the sgRNA and target DNA binding behavior and other characteristics that are relevant to the thermostability of a thermophilic Cas9.

## Results and discussions

### Overall structure of the binary complex of GeoCas9 and sgRNA

To solve the structure of ribonucleoprotein complex of GeoCas9, the purified protein of GeoCas9 that contains the co-expressed sgRNA was subjected to cryo-EM analyses. We also constructed a variant that harbors inactivated HNH domain, aiming to solve the complex with the target dsDNA. For this purpose, residue His582, presumed as the catalytic residue of HNH domain based on sequence alignment of GeoCas9 and homologous Cas9, was substituted by Ala (Extended Data Fig. 1) ^11,12,31^. The variant H582A yielded higher resolution structure than the WT protein (data not shown), thus we decided to refine and analyze the structure of GeoCas9^H582A^/sgRNA in the following content.

The cryo-EM structure of the GeoCas9^H582A^/sgRNA binary complex was solved at 3.2 Å resolutions (Extended Data Table 1). GeoCas9 adopts a bilobed architecture, which comprises NUC lobe and REC lobe that are connected by the arginine-rich bridge helix (BH) (Fig. 1a and b). The NUC lobe comprises RuvC, HNH, wedge (WED) and PAM interacting (PI) domains (Fig. 1b). As do other type C Cas9s, the former two domains should account for the cleavage of target DNA while the C-terminal WED and PI domains shall involve in the binding of sgRNA^12,26^. The REC lobe of GeoCas9 comprises REC1 and REC2 and is smaller than that of the well-studied SpyCas9, which contains an additional REC2 domain between REC1 and REC3 (Extended Data Fig. 2). It has been demonstrated that removing REC2 shows minimal influences on the SpyCas9 activity^7^. These results, together with the fact that several Cas9s also harbor small REC lobe as does GeoCas9^11,13,14^, showcase a minimal requirement for a Cas9 REC lobe.

**Fig. 1.**
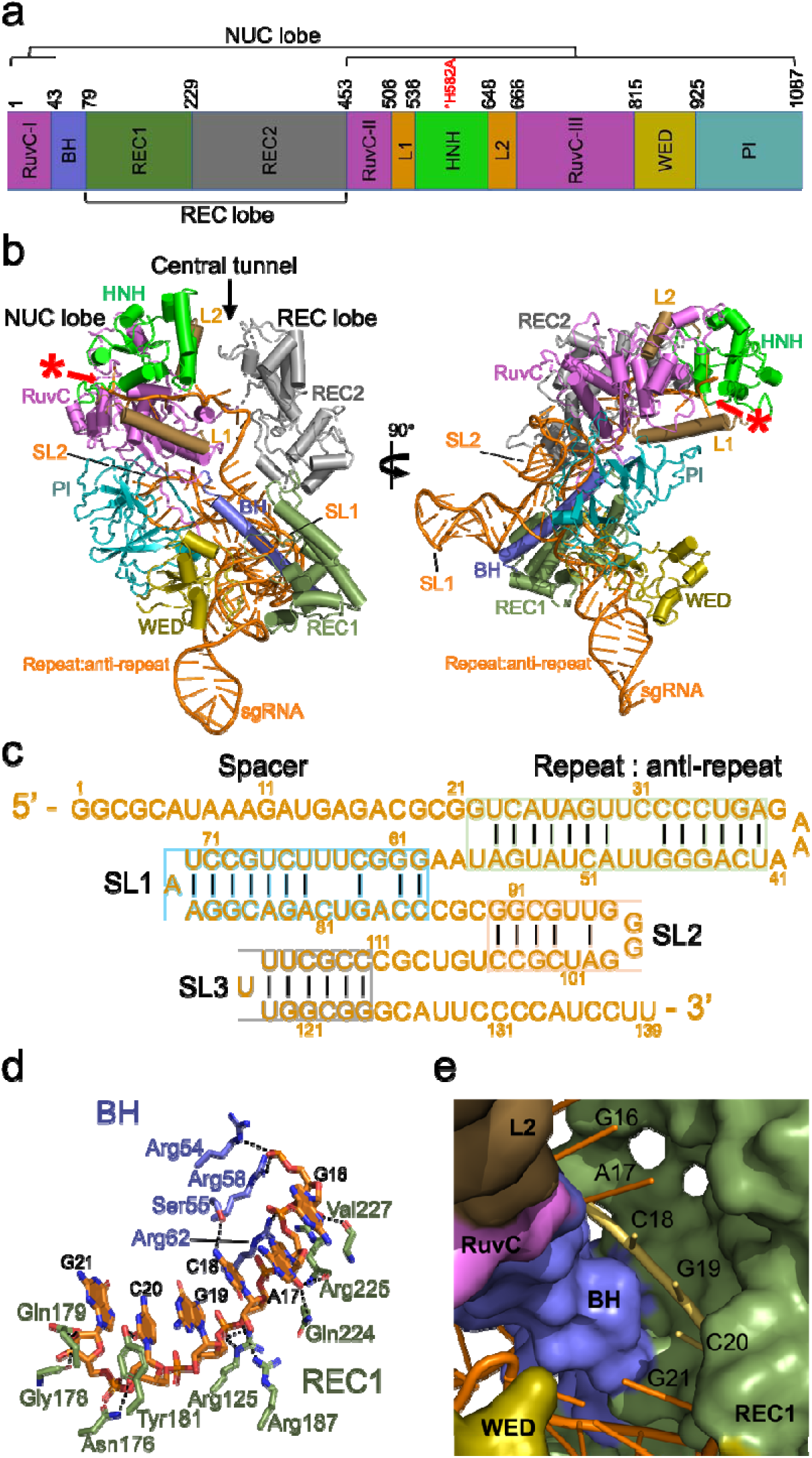
The GeoCas9^H582A^/sgRNA binary complex structure. **a**, The domain alignment of the HNH-inactive variant GeoCas9^H582A^. The numbers shown above the drawing are the amino acid numberings. **b**, The GeoCas9^H582A^/sgRNA complex structure (PDB ID, 8JTR) presented in a cartoon model with each domain and sgRNA segment labeled. Two views are related by 90° at Y-axis. The L1-crevice architecture is indicated by the red asterisk. **c**, The sequence and sub-structure of sgRNA in the binary structure. **d**, The amino acid interaction networks of nt G16-G21 of sgRNA in the binary complex. The dashed line indicate distance < 3.5 Å. **e**, The nt G16-G21 of sgRNA and the surrounding GeoCas9 domains are displayed in cartoon and surface models, respectively. The putative nucleation site comprising nt C18-C20 of the sgRNA is shown in yellow color. SL, stem loop.

The sgRNA that comprises spacer (nt G1-G21), repeat (nt G22-A37) and tracrRNA (nt U42-U139) was presumed to fold into repeat:anti-repeat region (nt G22-U57) and three stem loops (SL, Fig. 1c)^26^. These sub-structures can be identified in our binary complex except for the SL3 (Fig. 1b). The most extensive RNA-protein interactions occur in the repeat:anti-repeat region, which is located in the lower part of the central tunnel formed by REC1, WED and PI domains (Fig. 1b and Extended Data Fig. 3). The SL1 forms few interactions to the BH helix and extends away from the REC lobe while the SL2 leans against RuvC and PI domain (Fig. 1b and Extended Data Fig. 3).

The electron density of the spacer region is incomplete such that only nt G1-C5 and nt G16-G21 can be clearly identified (Extended Data Fig. 4). The former part is located in a slit, termed L1-crevice, formed by L1 helix, HNH and RuvC domain, while the latter is located in a region formed between BH helix and REC1 domain (Fig. 1b). Compared with nt G1-C5 that is loosely bound in the L1-crevice and form few contacts to the protein, segment G16-G21 forms extensive interactions to a number of polar residues and exhibits an A form-like pre-ordering structure that is considered to facilitate the base pairing to the TS-DNA (Fig. 1d and Extended Data Fig. 3)^20^. As nt G16, A17 and G21 are embedded within BH helix and REC1 domain, nt C18-C20 are exposed to the bulk solvent (Fig. 1e). This suggests that nt C18-C20 might serve as the nucleation site and play a key role in the binding of TS-DNA, as proposed in previous reports^20,32^.

### Ternary structure of GeoCas9 in complex with target DNA

To reveal the DNA binding mode of GeoCas9, we prepared and solved the complex of GeoCas9^H582A^/sgRNA and the 29-bp target dsDNA at 3.1 Å resolutions (Extended Data Table 1). Despite a fully paired dsDNA was used, a partial dsDNA that contains nine nucleotides in the NTS-DNA strand was observed in the ternary structure (Fig. 2a and b). This should be a result of the NTS-DNA strand being cleaved by the RuvC domain while the TS-DNA remaining intact owing to the lack of HNH activity.

**Fig. 2.**
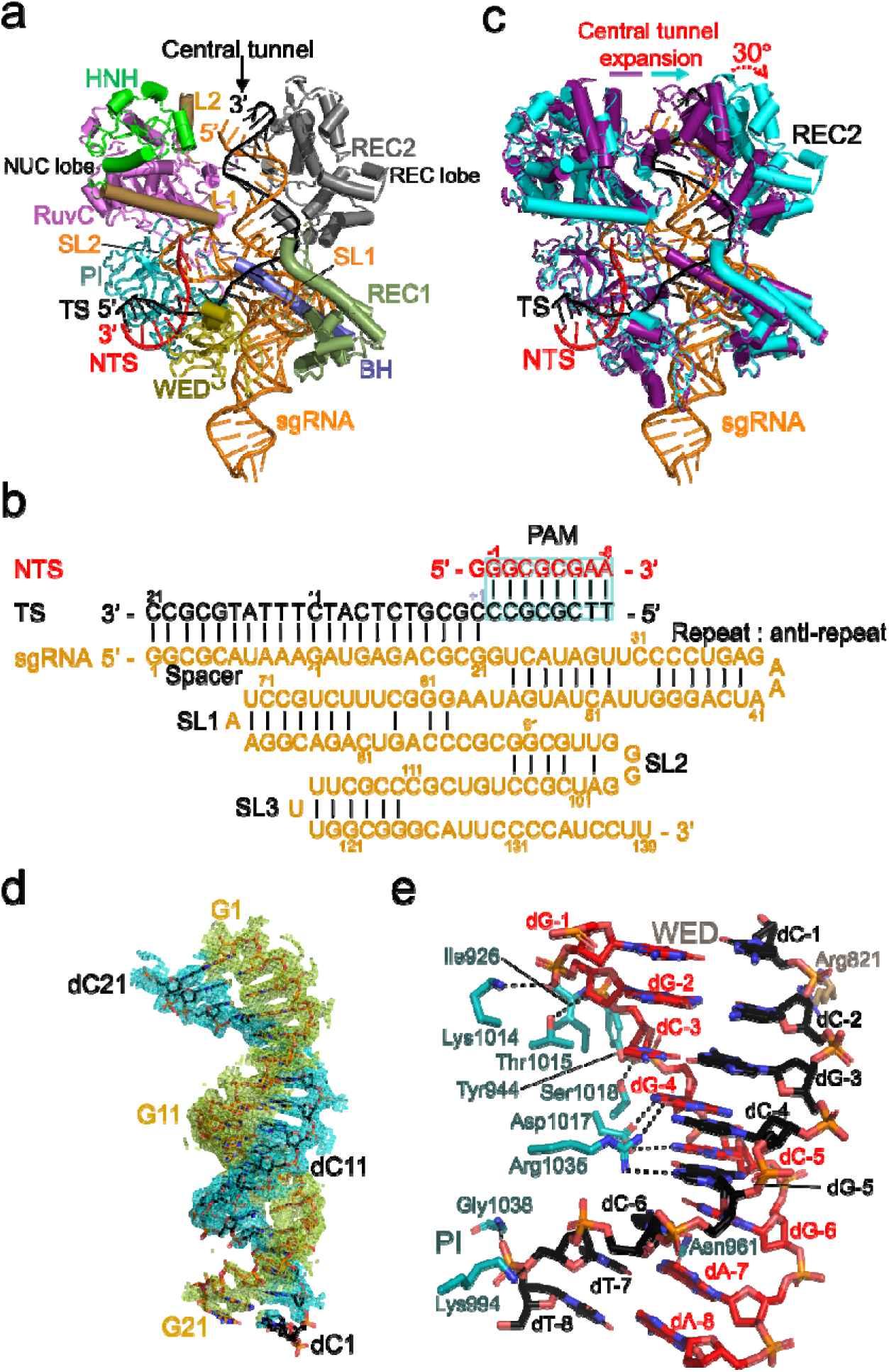
The GeoCas9^H582A^/sgRNA/dsDNA ternary complex structure. **a**, The GeoCas9^H582A^/sgRNA/dsDNA complex structure (PDB ID, 8JTJ) presented in a cartoon model with each domain, sgRNA segments, TS-DNA and NTS-DNA labeled. **b**, The sequence and sub-structure of sgRNA and partial dsDNA bound in the ternary structure. **c**, The binary (purple) and ternary (cyan) complex structure of GeoCas9^H582A^ are superimposed. Note that the central tunnel is expanded in the ternary complex. The REC2 domain in the ternary complex is tilted by 30° relative to that in the binary complex. **d**, The electron density maps of RNA:DNA duplex observed in the central tunnel. **e**, The GeoCas9 and PAM interaction network. The TS-, NTS-DNA and protein residues are displayed in black, red, sand and cyan sticks. The dashed line indicate distance < 3.5 Å.

Aligning the binary and ternary structures yields a root-mean-square deviation value of 1.4 Å for 758 equivalent Cα atom, suggesting that the two complexes share a similar overall architecture. The most notable variation is observed in the REC2 domain, which deviates outwards from the central tunnel by ∼30° (Fig. 2c). The movement of REC2 domain leads to the expansion of the central tunnel that serves to house the RNA:DNA duplex (Fig. 2c and Extended Data Fig.5), a phenomenon that has been reported in other Cas9^12,20^. The electron density of the entire spacer region (nt G1-G21) of the sgRNA can be clearly identified in the ternary complex, which should be owing to the stabilization effects conferred by the base pairing of the TS-DNA (Fig. 2d). The RNA:DNA duplex forms interactions to BH helix, REC1 and REC2 domains (Extended Data Fig. 6), while the PAM duplex is sandwiched by PI and WED domain (Extended Data Fig. 7). The GeoCas9 and PAM interaction occur in both DNA strands, that nt dG-5 of TS-DNA and nt dG-4, dC-5 and dA-7 of NTS-DNA form hydrogen bonds to residue Arg1035, Asp1017 and Asn961, and residue Arg821, Ile926, Tyr944, Lys994, Lys1014, Thr1015, Ser1018 and Gly1038 form polar interactions with the backbone of the PAM (Fig. 2e).

It is well acknowledged that the HNH domain along with the L1 helix would undergo dramatic conformational change to cleave the TS-DNA (Extended Data Fig. 8a)^6,12^. Nonetheless, the HNH domain and L1 helix in GeoCas9^H582A^/sgRNA/dsDNA complex only slightly deviated from their positions in the binary complex (Fig. 2c and Extended Data Fig. 8b). To be specific, the L1 helix is tilted by 20° in the ternary complex (Extended Data Fig. 9). This phenomenon is similar to the ternary structure of Cas9 from *S. aureus*, whose HNH domain is not transitioned to the DNA cleavage site (Extended Data Fig. 8c)^14^. Although the structure of SaCas9/sgRNA has been reported recently, the true conformation of sgRNA-bound status remains uncertain as this structure contains an inhibitor bound to the PAM site (Extended Data Fig. 8c). These results indicate that the GeoCas9^H582A^/sgRNA/dsDNA reported here does not exhibit a catalytic status when HNH domain approaches TS-DNA. Instead, it is suspected that we may have captured a transition status when the HNH and L1 helix become mobile and are readily to conduct conformational change.

### The “L1-crevice” serves as a sgRNA storage site

The observations including the presence of 5’ spacer in the L1-crevice in the binary complex attracted our attention (Fig. 1b and 3a). It has been demonstrated that the 10 nucleotides on the most 5’-part of crRNA is critical in forming the RNA:DNA duplex and are indispensable for the DNA cleavage activity^7,33,34^. But the location of this 5’ segment in Cas9 ribonucleoprotein complex remains elusive. From inspecting all deposited Cas9/sgRNA complex structures, we found that the binary complexes of SpyCas9 and Nme1Cas9 contain unmodeled electron densities in the L1-crevice corresponding region (Extended Data Fig. 10a)^12,20^. A recent report demonstrates that HpaCas9/sgRNA structure in complex with a broad-spectrum inhibitor contains a short fragment of RNA in the L1-crevice (Extended Data Fig. 10b), though no further discussions on this event have been provided^11^. According to these findings, the residence of 5’-end spacer in the L1-crevice could be a universal phenomenon of Cas9. In addition, we suspect that the insertion of the 5’ spacer could also contribute to the construction of the L1-crevice. Therefore, the L1-crevice might collapse upon the binding of target DNA when the loosely associated 5’ RNA fragment is extracted to form the RNA:DNA duplex. Subsequently, the L1 helix and the HNH domain shall be able to translocate to approach the TS-DNA cleavage site.

**Fig. 3.**
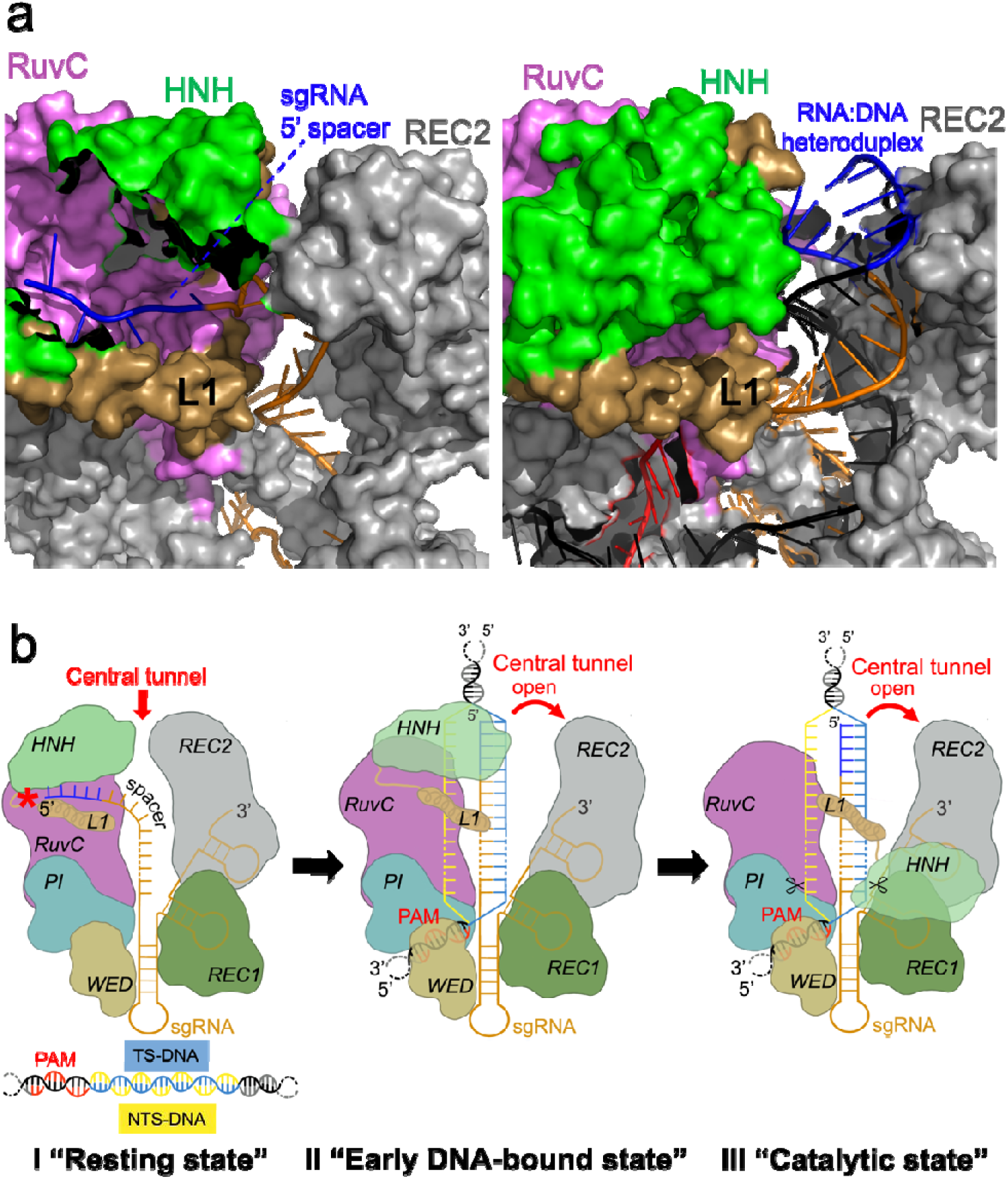
The L1-crevice. **a**, The L1-crevice formation in GeoCas9^H582A^/sgRNA (left panel) and collapse in GeoCas9^H582A^/sgRNA/dsDNA complex (right panel). The 5’ spacer that accommodates in the L1-crevice is colored in blue. **b**, A proposed process of Cas9 action. I, the sgRNA and Cas9 assemble to afford a resting binary complex, in which the 5’ spacer is stored in the L1-crevice (noted by the red asterisk) form by HNH, RuvC domain and L1 helix. II, the target dsDNA binds via the PAM and unwinds. The TS-DNA pairs with the sgRNA via the spacer region to yield a RNA:DNA duplex that causes the expansion of the central tunnel. The withdrawal of 5’ spacer from the L1-crevice leads to the destabilization of HNH domain and L1 helix. III, the HNH domain translocates to carry out DNA cleavage.

Based on these results, we presumed that the L1-crevice should be a transient storage site of 5’ spacer in the resting state, which might play a role in Cas9 activation. First, the sgRNA and Cas9 assemble to construct the binary complex as a resting state (I, Fig. 3b). As nt 18-20 in GeoCas9 is exposed and ready for the binding of target DNA, the upstream part (nt 1-13 in GeoCas9) stretches across the central tunnel with the most 5’ region (nt 1-5 in GeoCas9) located in the L1-crevice. Then, the target dsDNA binds via the PAM recognition site, leading to a stage termed DNA-bound state (II, Fig. 3b). The PAM proximal region is unwound to afford TS-DNA that pairs with spacer of the sgRNA. The 5’ spacer of sgRNA is withdrawn from the L1-crevice to construct the RNA:DNA duplex, which pushes the REC2 domain outward to expand the central tunnel and render the complex an open conformation. The absence of the 5’ spacer of sgRNA results in the collapse of the L1-crevice, which allows L1 helix and HNH domain to undergo the subsequent conformational change. Finally, the relocated HNH domain approaches and cleaves the TS-DNA (III, Fig. 3b).

### Molecular mechanism underlying thermostability of GeoCas9

Next, we attempted to explore the molecular mechanism underlying the thermostability of GeoCas9. The most well acknowledged protein thermostability-associated features include disulfide linkage and higher proportion of proline^35–37^. We found that GeoCas9, as well as all reported Cas9 structures, lack disulfide bond. As Cas9 action involves dramatic conformation change, it is reasonable that the protein should maintain flexibility and is not constrained by intramolecular covalent linkage. Notably, we found that GeoCas9 contains 4.3% proline, higher than the mesophilic SpyCas9 (Fig. 4a and Extended Data Table 2). Intriguingly, another thermophilic Cas9 from *A. celllulolyticus* (AceCas9) that exhibits optimal activity at 50 °C contains 5.6% Pro (Fig. 4a and Extended Data Table 2). Pro residues are distributed across all domains in GeoCas9 except for L1 and L2 helix that undergo significant conformational change during DNA cleavage (Fig. 4b and Extended Data Table 3). There are three Cas9s including Nme1Cas9, CdiCas9 and AnaCas9 that also contain around 4 – 5% Pro, whereas their activity has only been assessed at 37 □ (Extended Data Table 2).^12,17,25^ It would be interesting to probe their thermal profiles to see if these Cas9s can operate under elevated temperature.

**Fig. 4.**
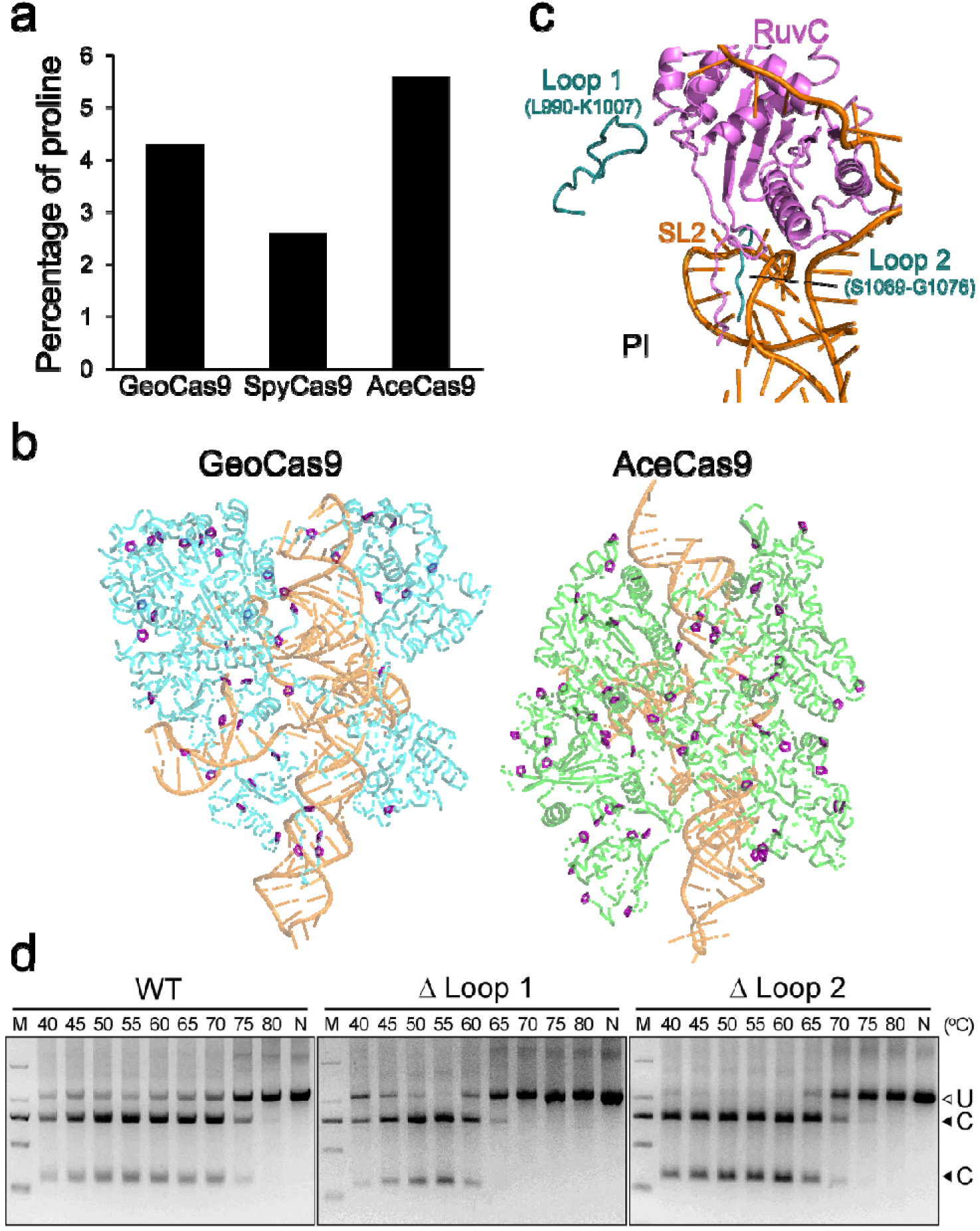
The thermostability-related feature of GeoCas9. **a**, The percentage of proline in total amino acid composition of Cas9 proteins whose thermal profiles have been demonstrated. **b**, The distribution of proline in the thermostable GeoCas9 and AceCas9. The overall structures of GeoCas9^H582A^/sgRNA/dsRNA (cyan; PDB ID, 8JTJ) and AceCas9/sgRNA/dsDNA (green; PDB ID, 8D2P) are displayed in cartoon models, with prolines shown in purple sticks. **c**, The PI domain in superimposed GeoCas9^H582A^/sgRNA (teal; PDB ID, 8JTR), Nme1Cas9/sgRNA (red; PDB ID, 6JDQ) and HpaCas9/sgRNA/AcrIIC4 (green; PDB ID, 8HNT) are displayed in cartoon models. Loop 1 and loop 2 of GeoCas9 are highlighted and indicated. The RuvC domain and partial sgRNA in GeoCas9/sgRNA binary complex are shown in magenta and orange models. **d**, The in vitro DNA cleavage activity of WT, loop 1- (Δ Loop 1) and loop 2-truncated (Δ Loop 2) variant GeoCa9 at indicated temperature. M, DNA marker; N, no enzyme control; U, uncut DNA fragment; C, cut DNA fragment.

In addition to the higher Pro ratio, we also noticed that GeoCas9 contains two additional loops in the PI domain in comparison with the closest related Nme1Cas9 and HpaCas9 (sequence identity vs. GeoCas9, 39 and 40%, respectively) (Extended Data Fig. 11). Loop 1 region comprising residue 990-1007 stretches to approach the RuvC domain, while the loop 2 region comprising residue 1069-1087 extends to the SL2 (Fig. 4c). To probe the role of these two loops in GeoCas9 thermostability, variants with these regions truncated were constructed and examined for the DNA cleavage activity. Compared with WT GeoCas9 that can cleave the substrate DNA at 75 □, the variant with loop 1 truncated showed very low activity at 65 □ and no activity above 70 □ (Fig. 4d). Deleting loop 2 also affects the thermostability of GeoCas9 but to a less degree, such that DNA cleavage activity can be detected at 70 □. These results clearly indicate a correlation between the loop 1 region and GeoCas9 thermostability. Despite loop 1 does not form substantial interactions to other domains, it covers a region comprising residue 1031-1036, whose counterpart in Nme1Cas9 and HpaCas9 is slightly longer and exposed to the bulk solvent (Extended Data Fig. 12). Therefore, we suspected that the unique PI domain architecture of GeoCas9 is a factor that contribute to the resilience of the protein at elevated temperature.

In conclusion, the cryo-EM structures of GeoCas9 in complex with sgRNA and target dsDNA have been resolved. Structural analyses indicate that L1-crevice is a transient storage site for the 5’ spacer segment of sgRNA, which could be a universal feature of Cas9. These findings lead to a proposition that the withdrawal of 5’ spacer from the L1-crevice could collapse the L1-crevice architecture, leading to the initiation of the HNH translocation. In addition, we also found that a higher proportion of Pro in total amino acid composition and a unique PI domain conformation are associated with the thermostability of GeoCas9. Altogether, our results provide structural and biochemical analyses to illustrate the molecular mechanism that could contribute to Cas9 thermostability, which increase our understanding of Cas9 biology and could be of interests to researchers that aim to increase the application potentials of Cas9-mediated gene editing.

## Acknowledgement

This work was supported by the National Key Research and Development Program of China (2022YFE0135300), Hubei Hongshan Laboratory (2022hszd030), the National Natural Science Foundation of China (32271318, 82341210 and 82103994); the Natural Science Foundation of Hubei Province (2022CFB360).

## Author contributions

P.S., X.L. and J.W.H. collected and analyzed data; P.S. and B.L. purified proteins, constructed variants and conducted biochemical assays; L.Z., J.M., C.C.C., and R.T.G. designed experiments; C.C.C. and R.T.G. conceptualized the project and wrote the manuscript.

## Materials and Methods

### Plasmid construction and mutagenesis

The *G. stearothermophilus* Cas9 (GeoCas9)-coding gene (GenBank no. WP_121625896.1) fused with a C-terminal 6 × His tag was cloned into a modified pET-32a that lacks TrxA tag. Variant GeoCas9-expressing plasmids were generated by using site-directed mutagenesis with mutagenesis oligonucleotides (Extended Data Table 4). The sgRNA-encoding gene, which was designated following the previous study^26^, was cloned on the C-terminus of WT or variant GeoCas9-coding gene. All plasmids were verified by direct sequencing.

### Expression and purification of GeoCas9/sgRNA binary complex

The recombinant plasmids that harbor GeoCas9-and sgRNA-expressing genes were transformed to *E. coli* BL21(DE3) and cultured in Luria-Bertani medium that contains 100 μg/mL ampicillin at 37 □. When cells grew to an OD600 ∼ 0.6, 0.5 mM isopropyl β-D-thiogalactopyranoside was supplemented for protein induction at 37 □ for 14 h. Cells were harvested by centrifugation at 7,000 rpm for 10 minutes and then resuspended in a lysis buffer containing 20 mM Tris-HCl [pH 7.5], 500 mM NaCl and 10 mM imidazole and disrupted by using French Press. The cell lysates were centrifuged at 17,000 rpm for 50 min at 4 □ to remove cell debris. The supernatant was then applied to a Ni-NTA column coupled on an AKTA Purifier system (GE Healthcare, Madison, WI) and target protein was eluted by 10-500 mM imidazole gradient. The target protein-containing fractions were pooled and dialyzed against a buffer containing 20 mM Tris-HCl [pH 7.5] buffer at 4 □ overnight and concentrated to 20 mg/mL. The protein solutions was loaded on to Superose 6 Increase 10/300 GL column (GE Healthcare, Madison, WI) that was equilibrated with a buffer containing 20 mM Tris-HCl [pH 7.5], 0.1 M KCl, 5 mM MgCl_2_, 0.5% glycerol (v/v) and 1 mM DTT. To prepare ternary complex, GeoCas9^H582A^/sgRNA binary complex was incubated with target dsDNA at a molar ratio of 1:1.25 at 37 □ for 30 min before loaded to the column. The fractions that contained target protein with expected molecular mass on the SDS-PAGE analysis were collected and concentrated to 20 mg/mL for further analysis.

### Preparation of target dsDNA

The 29-bp target dsDNA was constructed by using two oligonucleotides that were chemically synthesized (Genecreate Biological Co. Ltd., Wuhan, China):

Target strand: 5’ - TTCGCGCCCGCGTCTCATCTTTATGCGCC - 3’

Non-target strand: 5’ - GGCGCATAAAGATGAGACGCGGGCGCGAA - 3’

Two oligonucleotides that were dissolved in ddH_2_O to a final concentration of 200 mM were mixed by 1:1 ratio. The mixture was heated at 95 □ for 5 min and cooled to 25 □ by the rate of 1 □ /min.

### Cryo-EM grid preparation and data collection

For binary complex sample preparation, 4 μL of purified protein (2 mg/mL) was applied on Cu R1.2/1.3, 300 mesh holy carbon grids (Quantifoil, Jena, Germany) which had been glow discharged with 15 mA at 0.39 mBar for 1 min. For ternary complex sample, 4 μL of purified protein (2 mg/mL) was applied on Au-flat R1.2/1.3, 300 mesh holy gold grids (Electron Microscopy Sciences, Hatfield, PA) which had been glow discharged with 15 mA at 0.39 mBar for 15 s. The grids were blotted for 5 s at 4 □ with 100% humidity, and plunge frozen into liquid ethane cooled by liquid nitrogen using a Vitrobot Mark IV (ThermoFisher Scientific, Hillsboro, OR). High-quality grids were loaded onto a Titan Krios electron microscope operated at 300 kV for data collection. Image stacks were recorded with a Gatan K3 summit direct detector using the super-resolution counting mode at a nominal magnification of 105,000×, resulting in a calibrated pixel size of 0.425 Å. The software of EPU (ThermoFisher Scientific, Hillsboro, OR) was used for fully automated data collection. Each stack of 40 frames was exposed for 2.5 s with preset defocus values ranged from -1.0 to 2.4 μM and total dose for each stack was ∼ 52 e^-^/Å^2^.

### Cryo-EM image processing

All cryo-EM data were processed by using cryoSPARC v3.3.1^38^. Patched contrast transfer function parameters were estimated for each micrograph from the dose-weighted averages of all frames with default parameters. For GeoCas9^H582A^/sgRNA binary complex, a total of 2,115 micrographs were selected and 1,179,045 particles were automatically picked using the cryoSPARC Blob picker and extracted with a box size of 256 pixels. After two rounds of 2D classification, 603,510 particles were selected, followed by one round of ab initio reconstruction with 3 classes and C1 symmetry imposed. Particles in two selected subgroups were subjected to one round of heterogeneous refinement to remove bad particles. After one round of non-uniform and local refinement, a final dataset of 305,427 particles was reconstructed to produce a 3.21 Å resolution map.

For GeoCas9^H582A^/sgRNA/dsDNA ternary complex, a total of 6,647 micrographs were selected, 3,732,546 particles were automatically picked, and 970,712 particles were selected after two rounds of 2D classification following the abovementioned methods. Selected particles were subjected to one round of heterogeneous refinement to further remove bad particles. After one round of non-uniform, a final dataset of 389,973 particles was reconstructed to produce a 3.08 Å resolution map. The resolution of final maps was estimated by gold-standard Fourier shell correlation (FSC) using the 0.143 criterion.

### Cryo-EM model building and refinement

The initial model for the GeoCas9^H582A^/sgRNA binary complex was built based on a template model generated by AlphaFold2^39^, which served as a template to build the model of the GeoCas9^H582A^/sgRNA/dsDNA ternary complex. The sgRNA and dsDNA were built manually with COOT^40^. The models were refined by using ISOLDE^41^ and PHENIX^42^ sequentially. The statistics of the map reconstruction and model building are summarized in Extended Data Table 1. All structural figures were prepared using UCSF ChimeraX^43^ and PyMOL (Schrödinger, LLC, NY).

### DNA cleavage assay

The DNA cleavage activity of GeoCas9 was conducted following the previously reported protocol with some modifications^26^. The oligonucleotides containing protospacer and PAM sequences of the target dsDNA were introduced into pET-32a vector. A pair of oligonucleotides that can amplify a fragment containing the target dsDNA sequence through PCR reaction were used to produce the GeoCas9 substrate DNA (Extended Data Table 5). For DNA cleavage reaction, 390 nM of WT or variant GeoCas9/sgRNA complex and 480 ng substrate DNA were added in a buffer containing 20 mM Tris-HCl [pH 7.5], 100 mM KCl, 5 mM MgCl_2_, 1 mM DTT and 5% glycerol (v/v) and incubated at indicated temperature (45-80 □) for 30 min. The reactions were quenched by heating at 95 □ for 15 min and analyzed by running the 2% agarose gels to visualize the DNA fragments.

