## Extended Data Table 1˜5 and Extended Data Fig. 1˜ Fig. 12 for "Structure of *Geobacillus stearothermophilus* Cas9: insights into the catalytic process and thermostability of CRISPR-Cas9"

**Extended Data Table 1.** Cryo-EM data collection, refinement and validation statistics

|  | **GeoCas9^H582A^/sgRNA**  (EMD-36650) | **GeoCas9^H582A^/sgRNA/dsDNA**  (EMD-36646) |
| --- | --- | --- |
| **PDB** | 8JTR | 8JTJ |
| **Data collection and processing** | | |
| Magnification  Voltage (kV) | 105,000  300 | 105,000  300 |
| Detector  Electron exposure (e^-^/Å^2^) | Gatan K3  52 | Gatan K3  52 |
| Defocus range (μm) | -1.0 to -2.4 | -1.0 to -2.4 |
| Pixel size (Å) | 0.425 | 0.425 |
| Symmetry imposed | C1 | C1 |
| Initial particle images (no.)  Final particle images (no.) | 603,510  305,427 | 970,712  389,973 |
| FSC threshold | 0.143 | 0.143 |
| Map resolution (Å) | 3.21 | 3.08 |
| **Refinement** |  |  |
| Initial model used (PDB code) | 5CZZ | 5CZZ |
| Model resolution (Å) | 3.7 | 3.5 |
| FSC threshold | 0.5 | 0.5 |
| Map sharpening *B* factor (Å^2^) | -117.3 | -136.6 |
| **Model composition** |  |  |
| Non-hydrogen atom | 10957 | 11639 |
| Protein residue  Nucleotide | 1056  105 | 1042  145 |
| ***B* factors (Å^2^)** |  |  |
| Protein | 103.31 | 70.28 |
| Nucleotide | 117.01 | 105.40 |
| **R.m.s. deviations** |  |  |
| Bond lengths (Å) | 0.003 | 0.003 |
| Bond angles (°) | 0.631 | 0.712 |
| **Validation** |  |  |
| MolProbity score | 1.92 | 2.01 |
| Clashscore | 7.76 | 9.27 |
| Poor rotamers (%) | 0.11 | 0.00 |
| **Ramachandran plot** |  |  |
| Favored (%) | 91.71 | 91.11 |
| Allowed (%) | 7.81 | 8.4 |
| Outliers (%) | 0.48 | 0.49 |

**Extended Data Table 2.** Proline composition and thermal profile of several Cas9 proteins

|  | **Number of Pro/total amino acid (%)^1^** | **Thermal profile** | **Reference** |
| --- | --- | --- | --- |
| GeoCas9 | 47/1087 (4.3) | Opti. temp., 50-60℃, active at 75℃ | This study and Ref. 1 |
| SpyCas9 | 35/1368 (2.6) | Active at 35-45 ℃ | Ref. 1 |
| AceCas9 | 64/1138 (5.6) | Opti. temp., 50 ℃ | Ref. 2 |
| Nme1Cas9 | 44/1082 (4.0) | Assessed at 37 ℃^2^ | Ref. 3 |
| CdiCas9 | 46/1084 (4.2) | Assessed at 37 ℃^2^ | Ref. 4 |
| AnaCas9 | 52/1101 (4.7) | NA | NA |
| SauCas9 | 26/1053 (2.5) | Assessed at 37 ℃^2^ | Ref. 5 |
| SthCas9 | 26/1122 (2.3) | Assessed at 37 ℃^2^ | Ref. 6 |
| HpaCas9 | 36/1055 (3.4) | Assessed at 37 ℃^2^ | Ref. 7 |
| SmuCas9 | 38/1065 (3.7) | Assessed at 37 ℃^2^ | Ref. 8 |

^1^The numbers in the parentheses are the percentage of Pro in the total amino acid of each Cas9.

^2^The activity was assessed at 37 ℃ while the thermostability and optimal temperature are not reported.

NA, not available

**Extended Data Table 3.** The number of proline in each domain of GeoCas9 and some homologues

|  | **RuvC** | **BH** | **REC1** | **REC2** | **L1** | **HNH** | **L2** | **WED** | **PI** |
| --- | --- | --- | --- | --- | --- | --- | --- | --- | --- |
| GeoCas9 | 15 | 1 | 2 | 8 | 0 | 5 | 0 | 8 | 8 |
| AceCas9 | 11 | 0 | 10 | 15 | 0 | 6 | 0 | 6 | 16 |
| Nme1Cas9  HpaCas9 | 15  14 | 0  0 | 6  4 | 8  4 | 0  0 | 5  6 | 0  0 | 4  3 | 6  5 |

**Extended Data Table 4.** Mutagenesis oligonucleotides used in this study

| **Name** | **Sequence (5'-3')** |
| --- | --- |
| H582A-F | GAAGTTGACCACGTGATCCCGTA |
| Loop 1 truncated-F | TGTACCCGAACGATCTGATTCGTATCGAAGACGTTTTCGTGTACTAT |
| Loop 1 truncated-R | GCTATCGATGGTTTTATAGTACACGAAAACGTCTTCGATACGAATCAG |
| Loop 2 truncated-F | CTCGAGCACCACCACCACCACCACTGAGAT |
| Loop 2 truncated-R | GGTGGTGGTGCTCGAGCGCCAGGCCAACACGTTTTTCACCA |

**Extended Data Table 5.** The oligonucleotides used to amplify the substrate DNA of cleavage assay

| **Name** | **Sequence (5'-3')** |
| --- | --- |
| Substrate DNA-F | GGTATCCCGACTCTGCTGCTG |
| Substrate DNA-R | GGAAGAAAGCGAAAGGAGCGG |

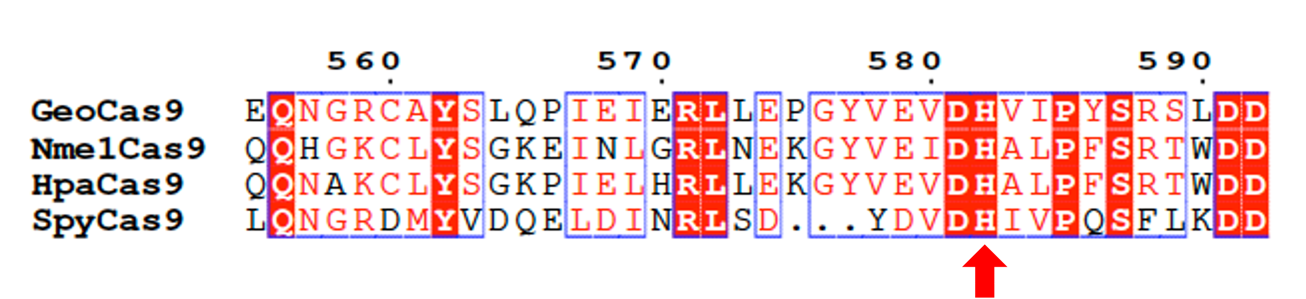

**Extended Data Fig. 1 Partial sequence alignment of GeoCas9 and some homologous Cas9.** The amino acid sequence of GeoCas9 (GenBank no., WP_121625896.1), Nme1Cas9 (GenBank no., WP_014574210.1), HpaCas9 (GenBank no., WP_005695805.1) and SpyCas9 (GenBank no., WP_010922251.1) are aligned by using ClustalOmega (https://www.ebi.ac.uk/jdispatcher/msa/clustalo). The catalytic residue H582 of HNH domain is indicated by the red arrow. The numbers on the top of the alignment are the amino acid numbering of GeoCas9. The figure are produced by Espript 3.0^9^.

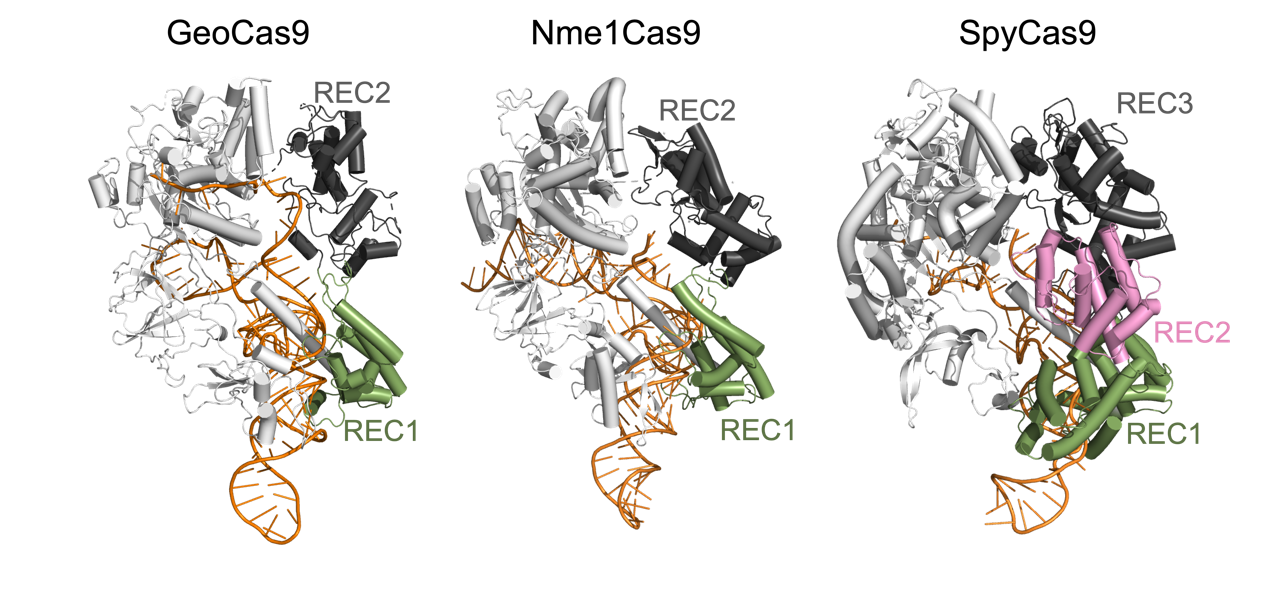

**Extended Data Fig. 2 REC lobe of GeoCas9, Nme1Cas9 and SpyCas9.** The structures of GeoCas9 (PDB ID, 8JTR), Nme1Cas9 (PDB ID, 6JDV) and SpyCas9 (PDB ID, 4ZT0) binary complex are displayed in cartoon models. The REC lobe-comprising domains are highlighted and labeled. Note that the REC2 in SpyCas9 is absent in the corresponding region in the other two Cas9 proteins.

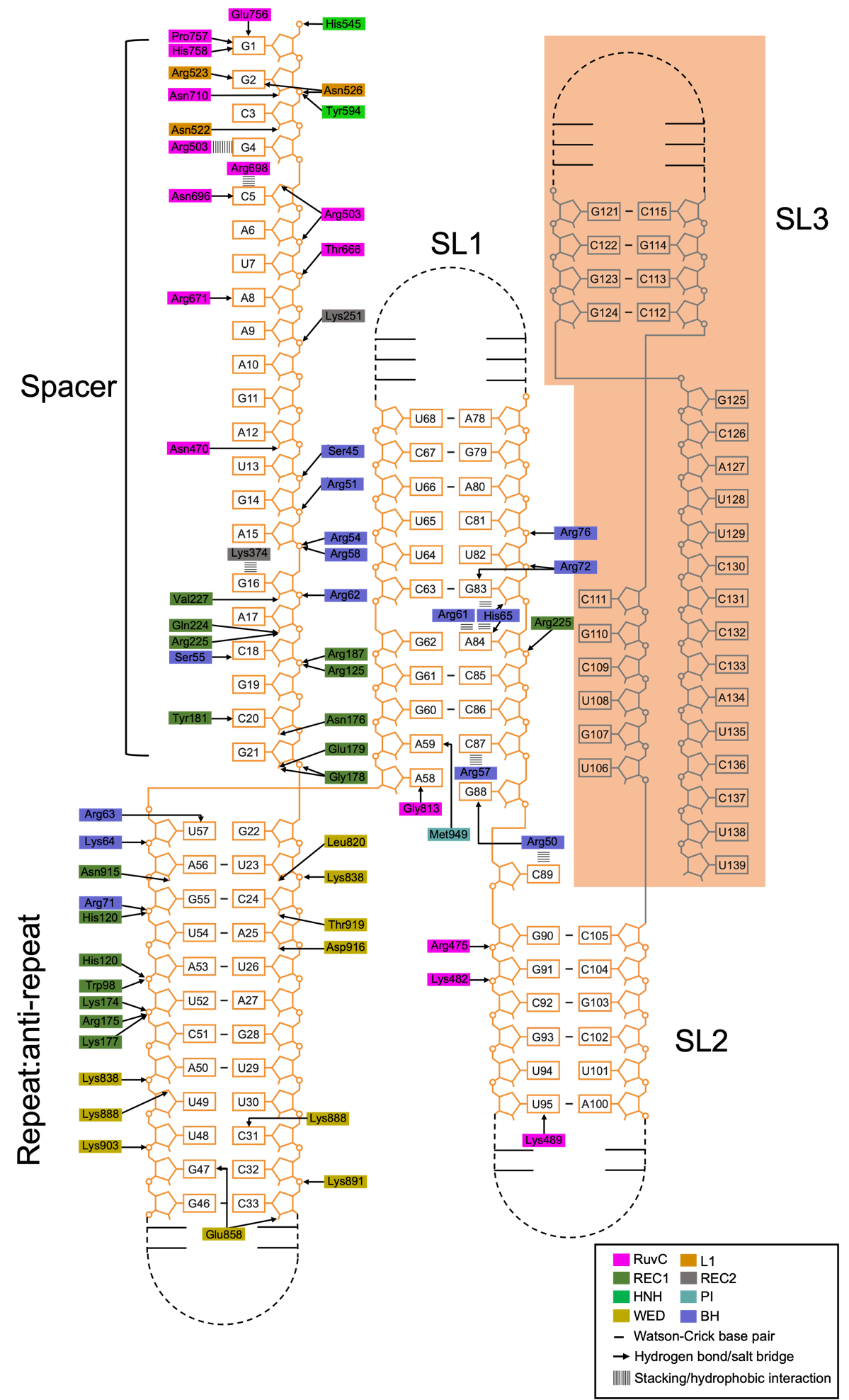

**Extended Data Fig. 3 The schematic presentation of amino acid-sgRNA interaction networks in GeoCas9^H582A^/sgRNA complex.**

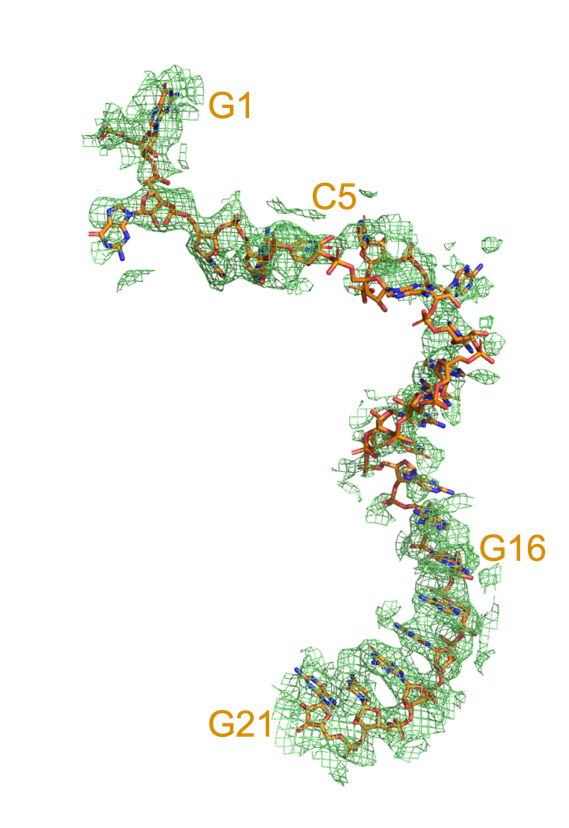

**Extended Data Fig. 4 The electron density map of sgRNA in GeoCas9^H582A^/sgRNA complex structure.** The electron density map and the sgRNA model are shown in mesh and sticks, respectively.

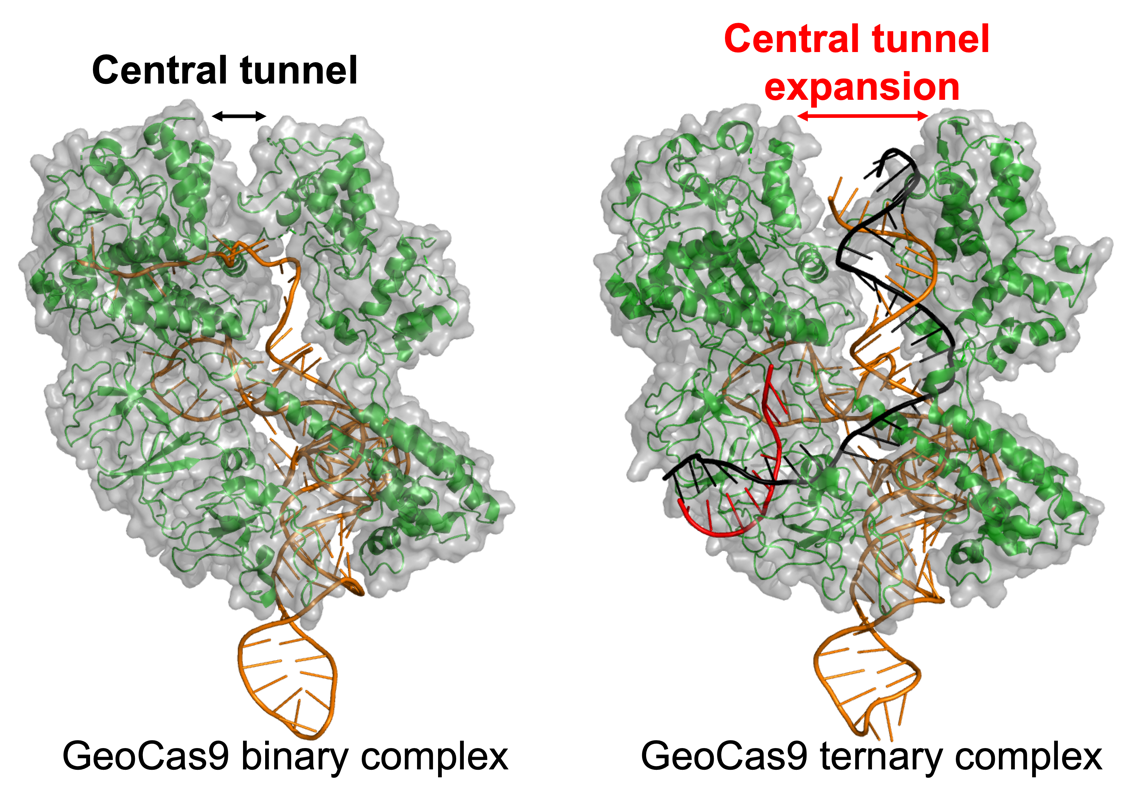

**Extended Data Fig. 5 Expansion of central tunnel in GeoCas9 upon the binding of target dsDNA.** The cartoon and surface model of binary and ternary complexes of GeoCas9 are displayed. sgRNA, TS-DNA and NTS-DNA are colored in orange, black and red, respectively.

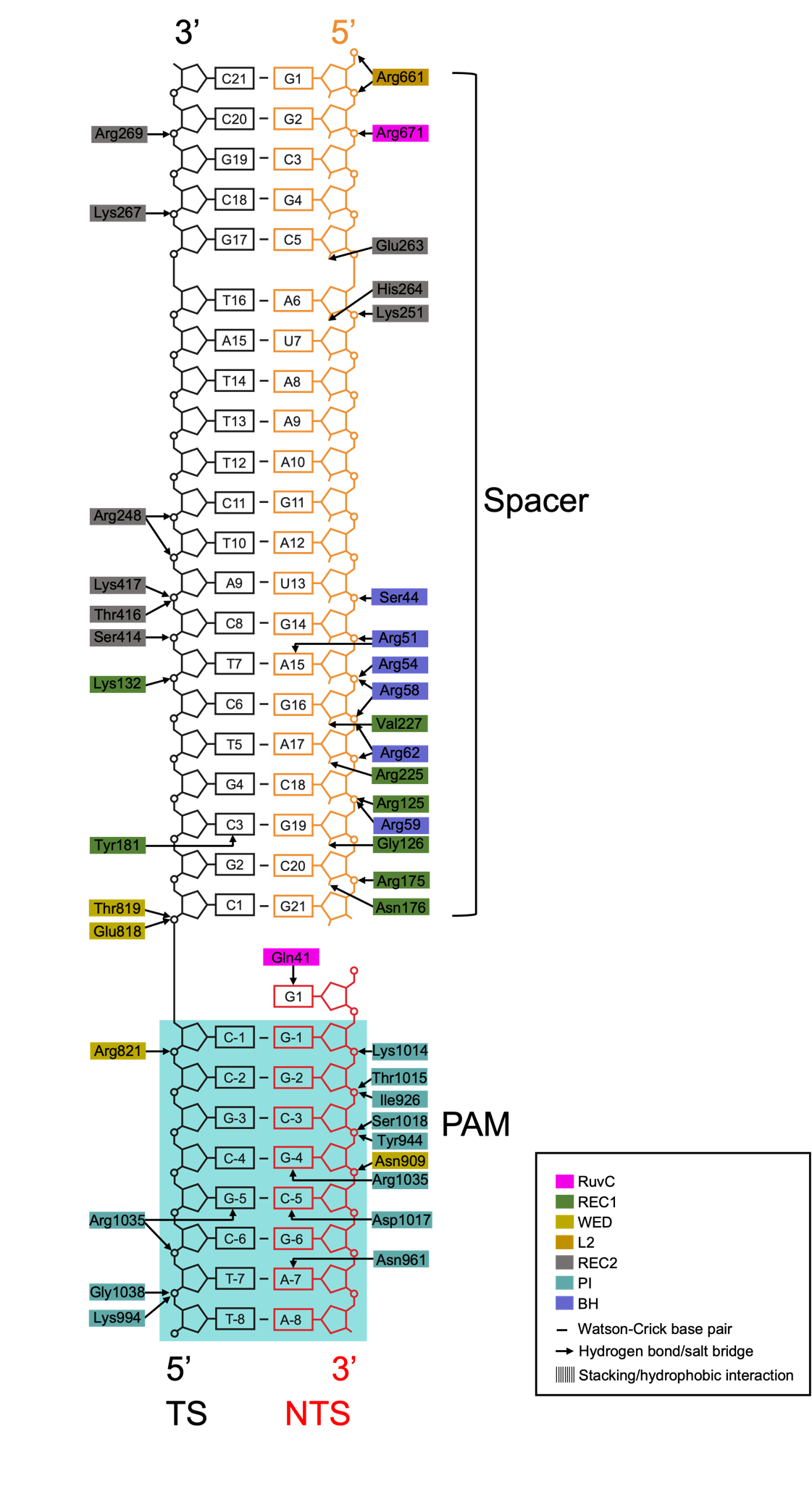

**Extended Data Fig. 6 The schematic presentation of amino acid-RNA:DNA duplex interaction networks in GeoCas9^H582A^/sgRNA/dsDNA complex.**

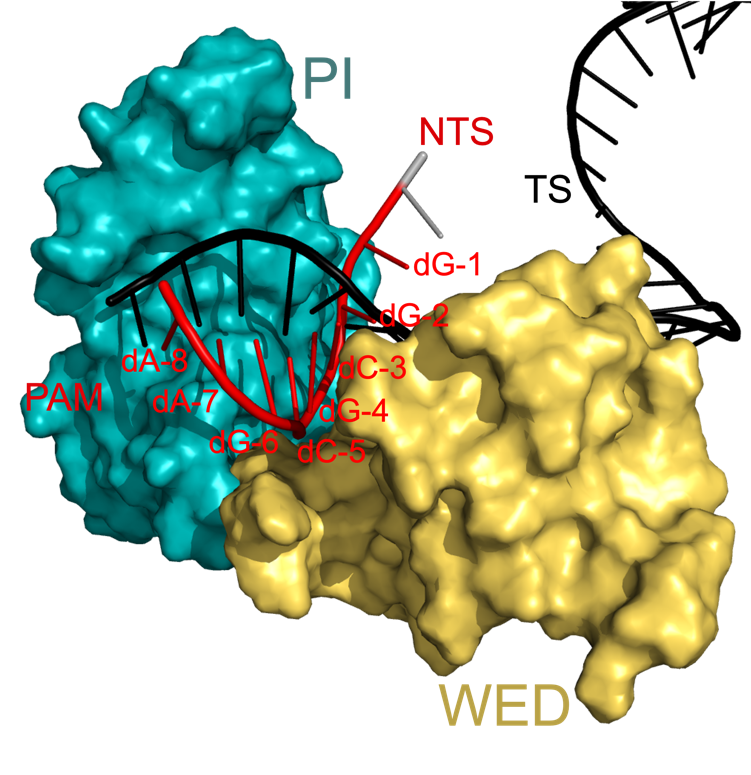

**Extended Data Fig. 7 The PAM-binding site in the GeoCas9^H582A^/sgRNA/dsDNA complex structure.** The PAM-binding site constituted by PI (teal) and WED (yellow) are displayed.

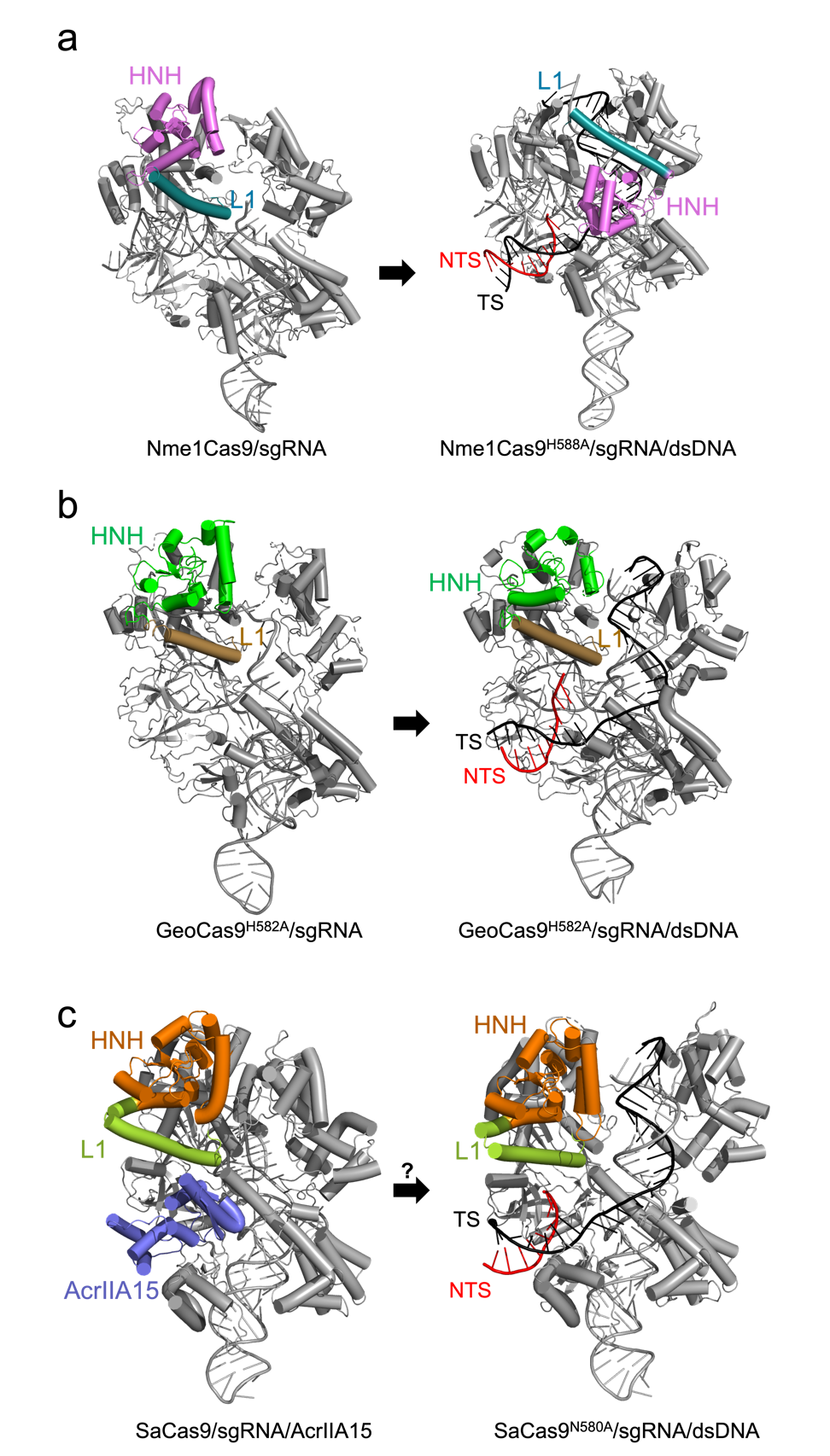

**Extended Data Fig. 8 The conformation change in reported Cas9 structures.** The cartoon model of **a**, Nme1Cas9 (binary complex, PDB ID, 6JDQ; ternary complex, PDB ID, 6JDV), **b**, GeoCas9 (binary complex, PDB ID, 8JTR; ternary complex, PDB ID, 8JTJ) and **c**, SaCas9 (binary and AcrIIA15 complex, PDB ID, 8JFT; ternary complex, PDB ID, 5CZZ) are displayed. The L1 helix and HNH domain in each structure are highlighted and labeled. Note that an inhibitor is bound in the SaCas9/sgRNA complex, thus the real conformation of the SaCas9/sgRNA complex remains uncertain.

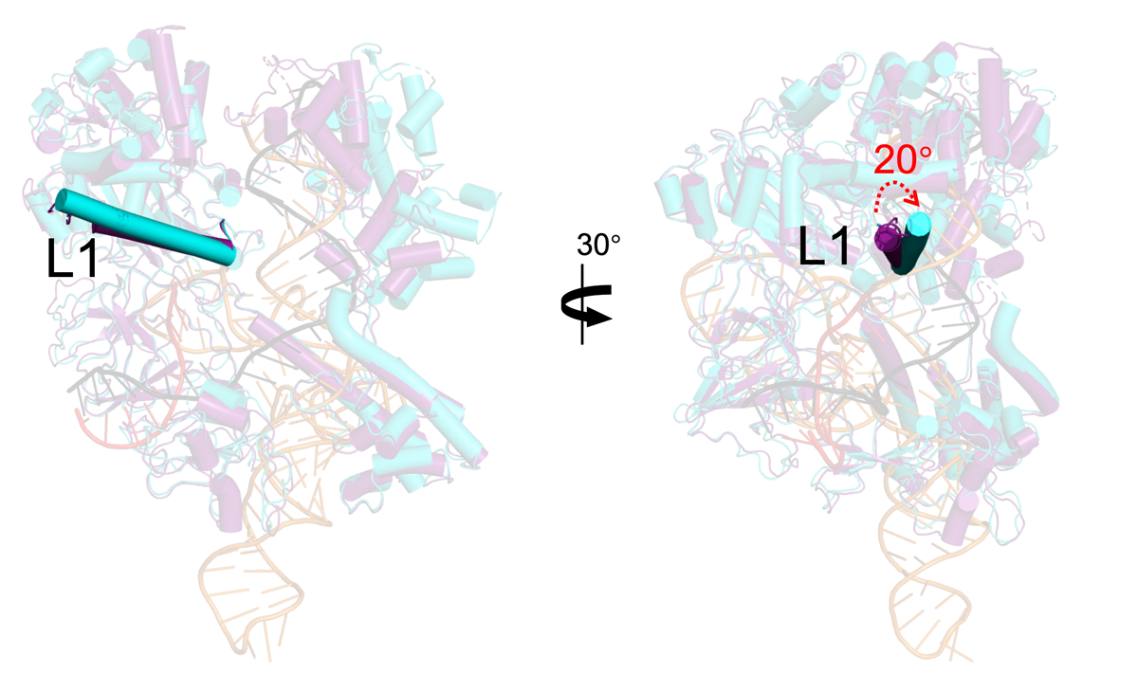

**Extended Data Fig. 9 The deviation of L1 helix of GeoCas9.** The structures of GeoCas9^H582A^/sgRNA (purple; PDB ID, 8JTR) and GeoCas9^H582A^/sgRNA/dsDNA (cyan; PDB ID, 8JTJ) displayed in cartoon models are superimposed. The sgRNA and target dsDNA in the ternary complex structure are shown as described in **Extended Data Fig. 5**. The L1 helix in both structures are highlighted and labeled. Two views are related by 30º at Y-axis.

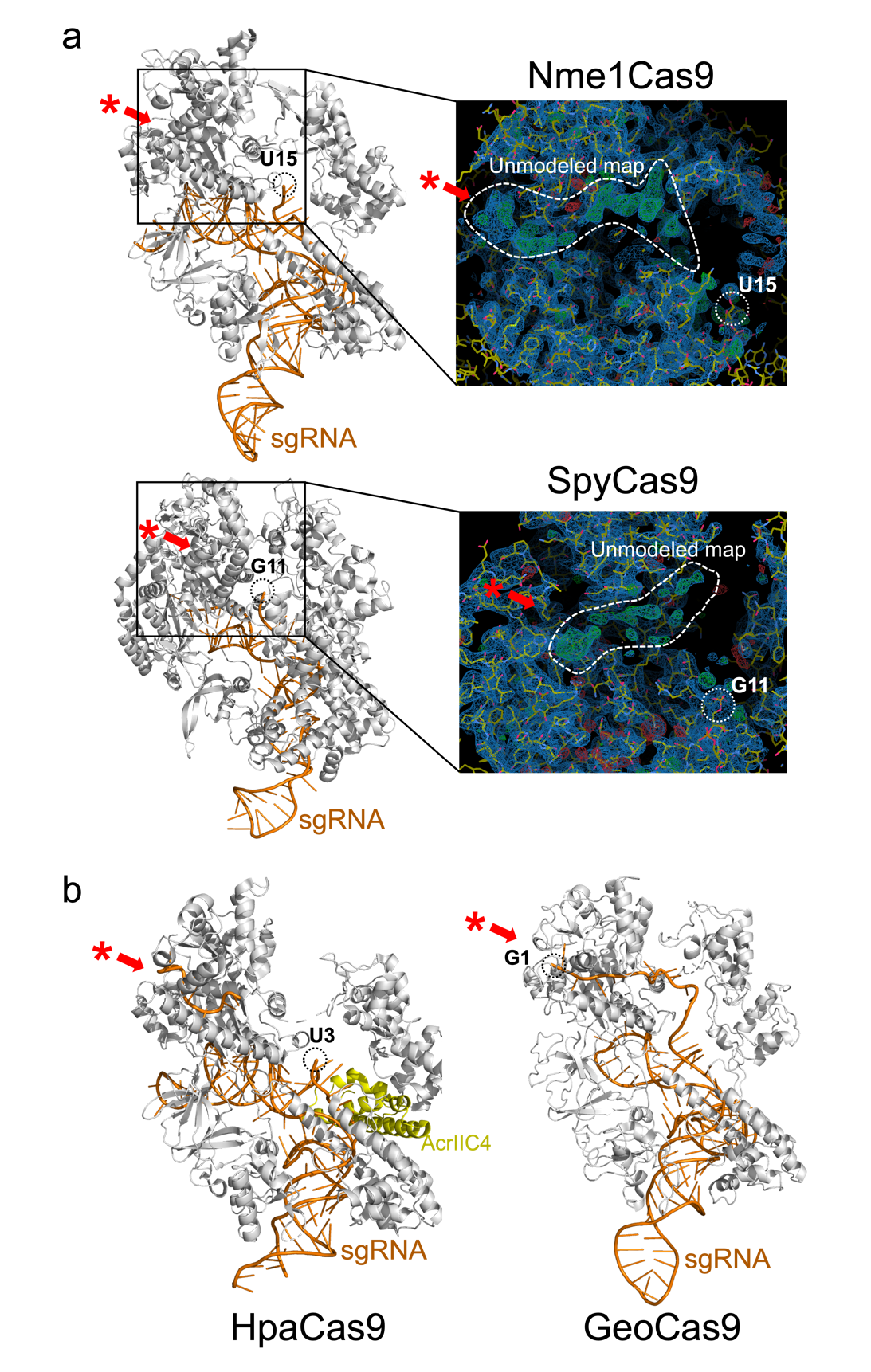

**Extended Data Fig. 10 The L1-crevice in Cas9. a**, The overall structure (left panel) and unmodeled electron density maps in L1-crevice corresponding region (right panel) of sgRNA binary complex of Nme1Cas9 (PDB ID, 6JDQ) and SpyCas9 (PDB ID, 4ZT0) are displayed. The unmolded maps are highlighted by dashed lines. **b**, The L1-crevice architecture in HpaCas9/sgRNA/AcrIIC4 (PDB ID, 8HNT) and GeoCas9/sgRNA (PDB ID, 8JTR) complex structures. The red asterisks indicate the location of L1-crevice, while the dashed circles indicate the most 5’-end nucleotide modeled in each structure.

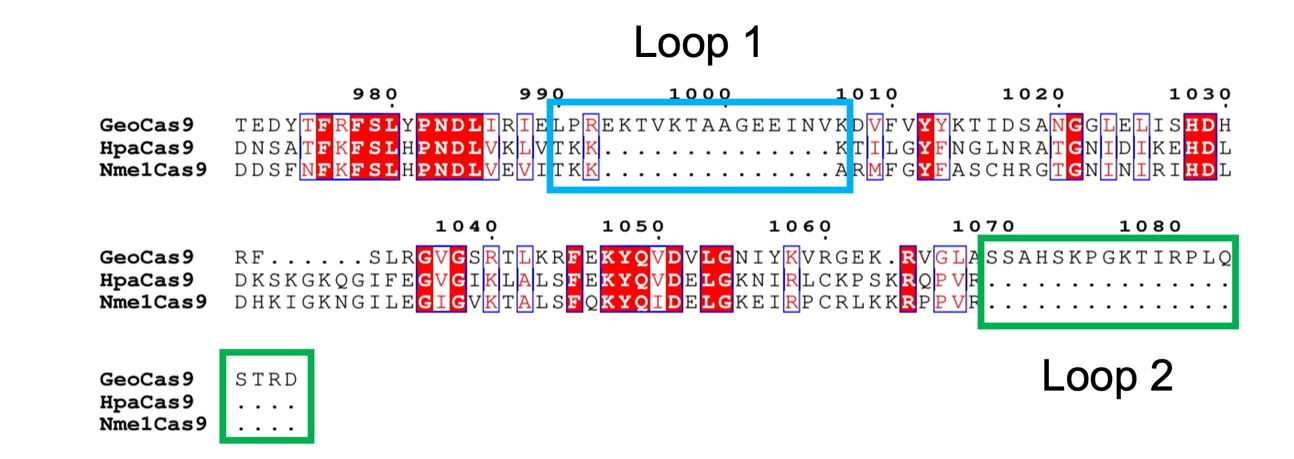

**Extended Data Fig. 11 The sequence alignment of PI domain in GeoCas9 and homologous Cas9 proteins.** The amino acid sequence of three Cas9 proteins are aligned and displayed as described in **Extended Data Fig. 1**, with two GeoCas9-unique loop regions framed and labeled.

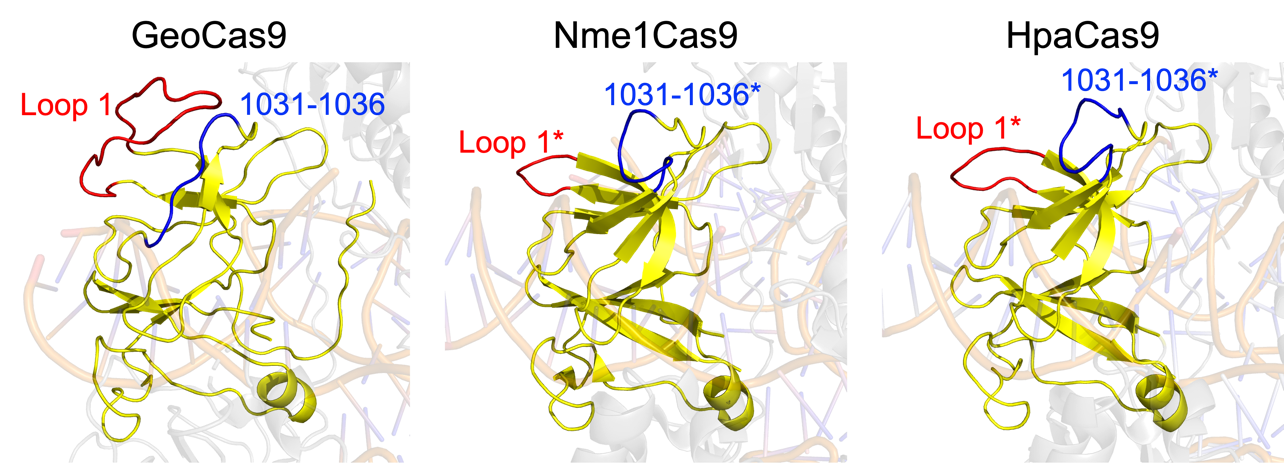

**Extended Data Fig. 12 Structural comparison of PI domain in GeoCas9 and homologues.** The cartoon models of PI domain in the ternary complexes of GeoCas9 (PDB ID, 8JTJ), Nme1Cas9 (PDB ID, 6JDV) and HpaCas9 (PDB ID, 8HNW) are displayed. The PI domains are shown in yellow, with loop 1 and segment 1031-1036 colored in red and blue, respectively. The corresponding regions of loop 1 and 1031-1036 in the other Cas9 are noted with asterisks.
